# Genome similarities between human-derived and mink-derived SARS-CoV-2 make mink a potential reservoir of the virus

**DOI:** 10.1101/2022.05.29.493871

**Authors:** Mohammad Khalid, Yousef Al-ebini

## Abstract

The SARS-CoV-2 has RNA as the genome, which makes the virus more prone to mutations. Occasionally, mutations help a virus to cross the species barrier. The SARS-CoV-2 infection to humans and minks (*Neovison vison*) are examples of zoonotic spillover. Many studies have been published on the analysis of human-derived SARS-CoV-2, here we performed mutation analysis on the minks-derived SARS-CoV-2 genome sequences. We analyzed all available full-length mink derived SARS-CoV-2 genome sequences on GISAID (214 from Netherlands and 133 from Denmark). We found that the mutation pattern in the Netherlands and Denmark derived samples were different. Out of a total of 201 mutations, we found in this study, only 13 mutations were common in the Netherlands and Denmark derived samples. We found 4 mutations prevailed in the Netherlands and Denmark mink derived samples and these 4 mutations are also reported to prevail in human-derived SARS-CoV-2.

## Introduction

The COVID-19 pandemic is caused by the SARS-CoV-2. The human transmission of the SARS-CoV-2 is a zoonotic spillover as consequence of mutational changes in the viral genome (1), these changes are ongoing process, and keep producing variants of the virus. The recent ΔH69/V70 variant of the virus reported to be more transmissible and deadly (2, 3). The ΔH69/V70 variant of SARS-CoV-2 has a deletion of two amino acids in spike (S) protein of the virus. The S protein is crown like structure and that’s how the virus got its name as the Coronavirus (4). The S protein of the virus uses the angiotensin converting enzyme 2 (ACE-2) as a receptor to enter into the host cells (4–6). Apparently; the ACE-2 receptor of different host cell, if has affinity towards the S protein of the virus, make the cell susceptible to viral entry and that makes the virus crosses the species barrier. This statement itself is very threatening and could have tremendous implications.

The minks (*Neovison vison*) in the Netherlands farms with symptoms of respiratory illnesses and higher mortality rate were found SARS-CoV-2 positive (7). The Munnink et al reported that people working in these farms were also infected with the SARS-CoV-2 (8) suggesting the virus can transmit back and forth between minks and human. Similar cases have been reported from the Denmark mink farms too (9–11), which suggests the infected minks, could be reservoir and/or potential source for the transmission of the virus to humans.

The European farmers are the highest producer of mink-fur form 6000 farms across the Europe. It accounts for 63% of the world’s production of mink-fur. Denmark is the leading producer of mink-fur and account for around 28% of world production (12).

The SARS-CoV-2 has a 29.9 kilobases, positive sense single-stranded RNA genome (+ssRNA) (13). The RNA makes the virus relatively more susceptible to mutational changes (14). Many reports have been already published about the mutation patterns in human derived SARS-CoV-2 (15–18) but we do not have sufficient information about the mutation patterns of the mink derived SARS-CoV-2 samples.

Our goal was to analyse the mink derived SARS-CoV-2 genome sequences and investigate the mutation pattern in the genome. In this study, we have examined all mink derived full-length SARS-CoV-2 genome sequences, available in GISAID (19) until November 4, 2020.

## Methods

### Mink Derived SARS-CoV-2 genome Sequences

In the present study, we analysed all available mink derived full-length SARS-CoV-2 genome sequences in GISAID (19). As on November 4, 2020, 347 sequences were available (set: November 2020), out of those, 214 sequences were from the Netherlands and 133 sequences were from mink farms in the Denmark. Further analysis for specific mutations, we used 755, SARS-CoV-2 genome sequences, available on January 22, 2021 (set: January 2021). Out of these 755 SARS-CoV-2 genome sequences, 454 were from Denmark, 278 were from Netherlands, 12 were from Poland, 7 were from Lithuania, and 4 sequences were from Canada.

### Mutational Analysis of the Mink Derived SARS-CoV-2 Genome Sequences

To analyse the November 2020 set of data of mink derived full length SARS-CoV-2 genome sequences, we have followed the method described elsewhere (20), briefly, a software tool for fragmenting nucleotide sequences, Fragmentation Tool_Version-1.0, were used to fragment the genome sequences. The 3000 bases long SARS-CoV-2 genome fragments were possible to align with MultiAlin online tool (21) with the reference strain of the virus (accession number NC_045512.2) (22). Each fragment contained 100 bases long overlapping sequences, to assure the fragments did not miss part of the genome sequence. We considered every variation from reference strain as a mutation, if variations repeated at more than one instance. The mutations prevailed in the mink derived SARS-CoV-2 genome November 2020 set of data, we further analysed in January 2021 set of data for their occurrence.

## Results

### Pipeline for the mutation analysis

### The SARS-CoV-2 Hosts Range

Table 1 provides the details of the SARS-CoV-2 genome sequences derived from various infected hosts and submitted to GISAID. These sequences were available as on January 22, 2021. The mink (*Neovison vison*) seemed to be highly susceptible to the SARS-CoV-2 infection; mink derived sample contributed 755 genome sequences, shown in the Table 1. The SARS-CoV-2 also infected several other animals viz. cat, pangolin, dog, tiger, lion, bat monkey, and mouse as shown in the Table 1.

**Table 1:** The SARS-CoV-2 hosts.

| S. No. | Name | Sequences |
| --- | --- | --- |
| 1 | Human (*Homo sapiens*) | 3,240,911 |
| 2 | Mink (*Neovison vison*) | 994 |
| 3 | Cat (*Felis catus*) | 70 |
| 4 | Lion (*Panthera leo*) | 37 |
| 5 | Dog (*Canis lupus familiaris*) | 28 |
| 6 | Pangolin (*Manis javanica*) | 19 |
| 7 | Tiger (*Panthera tigris jacksoni*) | 13 |
| 8 | Otter (*Aonyx cinereus*) | 5 |
| 9 | Mouse (*Mus musculus*) | 4 |
| 10 | Bat (*Rhinolophus malayanus*) | 4 |
| 11 | Bat (*Rhinolophus shameli*) | 2 |
| 12 | Bat (*Rhinolophus affinis*) | 1 |
| 13 | Monkey (*Chlorocebus sabaeus*) | 1 |
| 14 | Pangolin (*Manis pentadactyla*) | 1 |

The SARS-CoV-2 complete genome sequences derived from various hosts, available in GISAID as on December 21, 2021.

### The Phylogenetic Tree of SARS-CoV-2 Genomes, Derived from Various Hosts

We wanted to analyse similarities between the SARS-CoV-2 genome sequences derived from various hosts, shown in Table 1. We used MEGAX software tool (23) to generate the phylogenetic tree with the genomic sequences available in GISAID. These sequences were derived from various hosts mentioned in Table 1. We used one SARS-CoV-2 genomic sequence derived from every host as a representative. The human derived SARS-CoV-2 genome sequence used as reference (NC_45512.2) to established the phylogenetic relationship between these sequences.

### The Mutations in SARS-CoV-2 genome, derived from Netherlands and Denmark Mink Farms

As shown in Table 1, the mink (*Neovison vison*) derived SARS-CoV-2 genome sequences were the second highest, indicating that mink is highly susceptible to the viral infection. We were curious to find mutation patterns in the mink derived SARS-CoV-2 genomes. We analysed 347 (November 2020 set; 214 from Netherlands + 133 from Denmark) SARS-CoV-2 genome sequences, available on November 4, 2020. We found 201 mutations in the SARS-CoV-2 genome. These mutations are occurring in every UTRs/ORFs of the mink derived SARS-CoV-2 genomes except the ORF-E (Figure 3, Supplementary Table 1). The viral sequences derived from Netherlands and Denmark mink farms behaved differently in the mutation patterns (Supplementary Table 1). Out of 201 mutations, only 12 mutations (C241T, T3037C, A5421G, ATA6510deletion, G11083T, C14408T, C15656T, A22920T, A23403G, C25936T, G26062T, and C26078T) were common among the Netherlands and Denmark derived samples. Interestingly, out of these, 5 mutations (C241T, T3037C, G11083T, C14408T, and A23403G) were also reported in human derived SARS-CoV-2 genomes (20).

**Figure 1:**
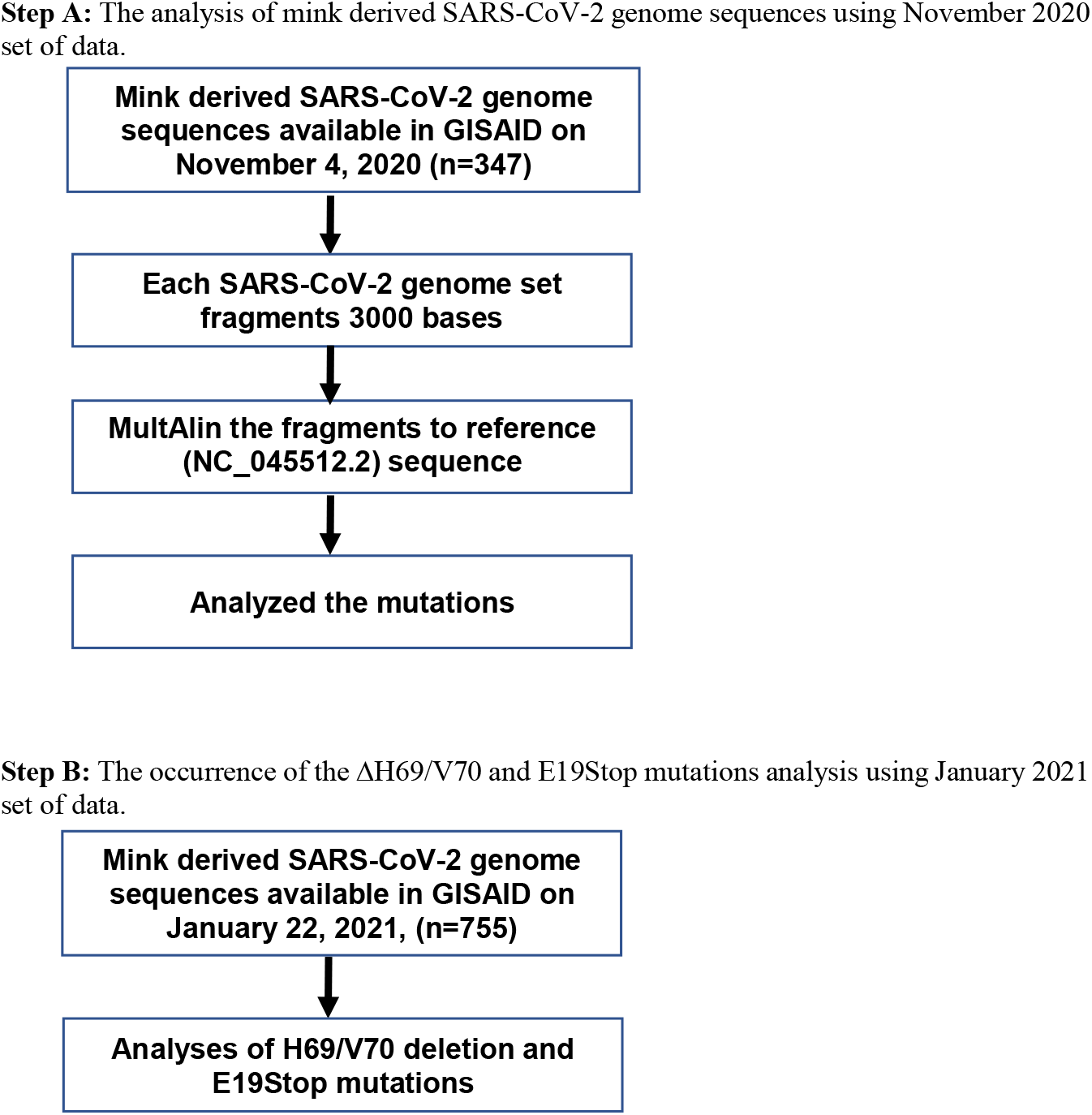
Flow chart of the Mink derived SARS-CoV-2 genome sequence analysis.

**Figure 2:**
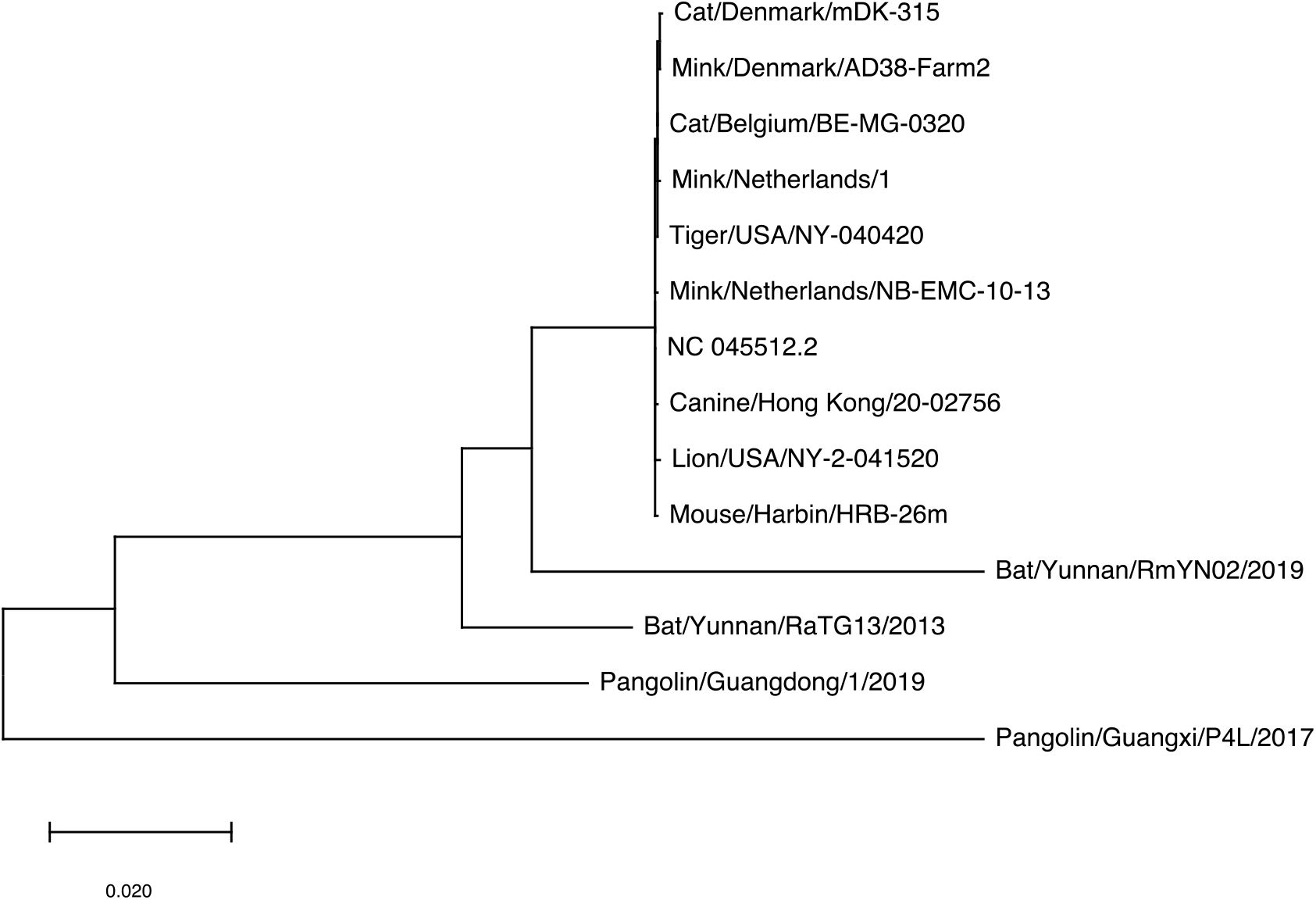
Phylogenetic Tree of the SARS-CoV-2 genomes, derived from various hosts.

**Figure 3:**
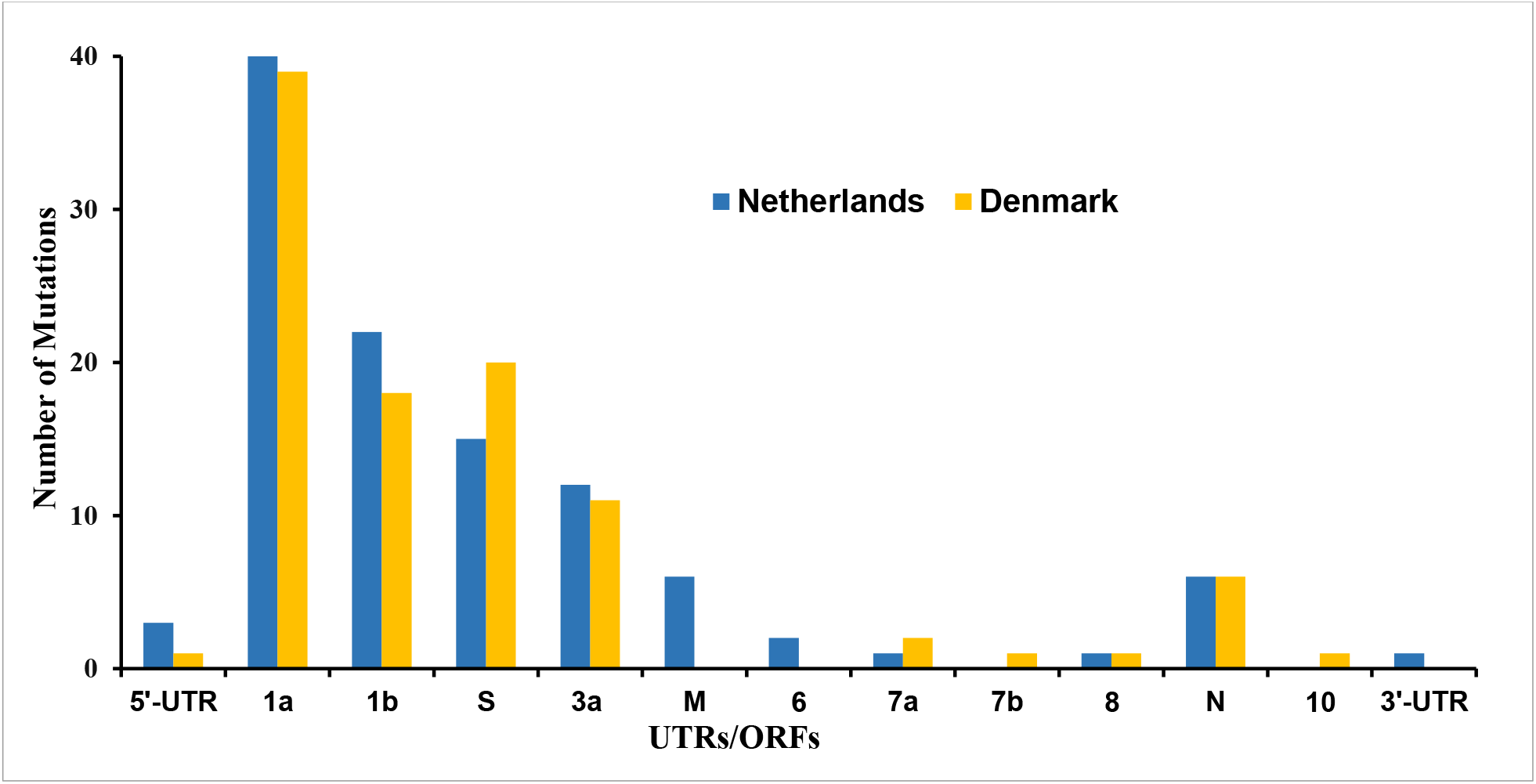
Mutations in various ORFs in the SARS-CoV-2 genome derived from the Netherlands and Denmark.

### The ΔH69/V70 and several other mutations prevailed in mink derived SARS-CoV-2 genome

We found total 201 mutations, out of those 18 mutations prevailed in the SARS-CoV-2 (occurred in more than 50% samples), Table 2 and Supplementary Table 1. Out of these18, 8 mutations (C241T, T3037C, ATA6510deletion, C14408T, C15656T, A22920T, A23403G, C25936T) were found prevailed in both Denmark’s as well as Netherland’s derived samples. The mutation ACATGT21766deletion, which translates to H69/V70 deletion in S protein, occurred in 84.2% samples derived from Denmark. None of the Netherlands’ sample were positive for ACATGT21766deletion (H69/V70 deletion) mutation, Table 2 and Supplementary Table 1. The A23403G mutation which translates to D614G mutation in the S protein became dominant in human derived SARS-CoV-2 (24) were also found prevailing in the Denmark’s (100%) and Netherlands’ (89.3%) mink derived SARS-CoV-2, Table 2, and Supplementary Table 1.

**Table 2:**
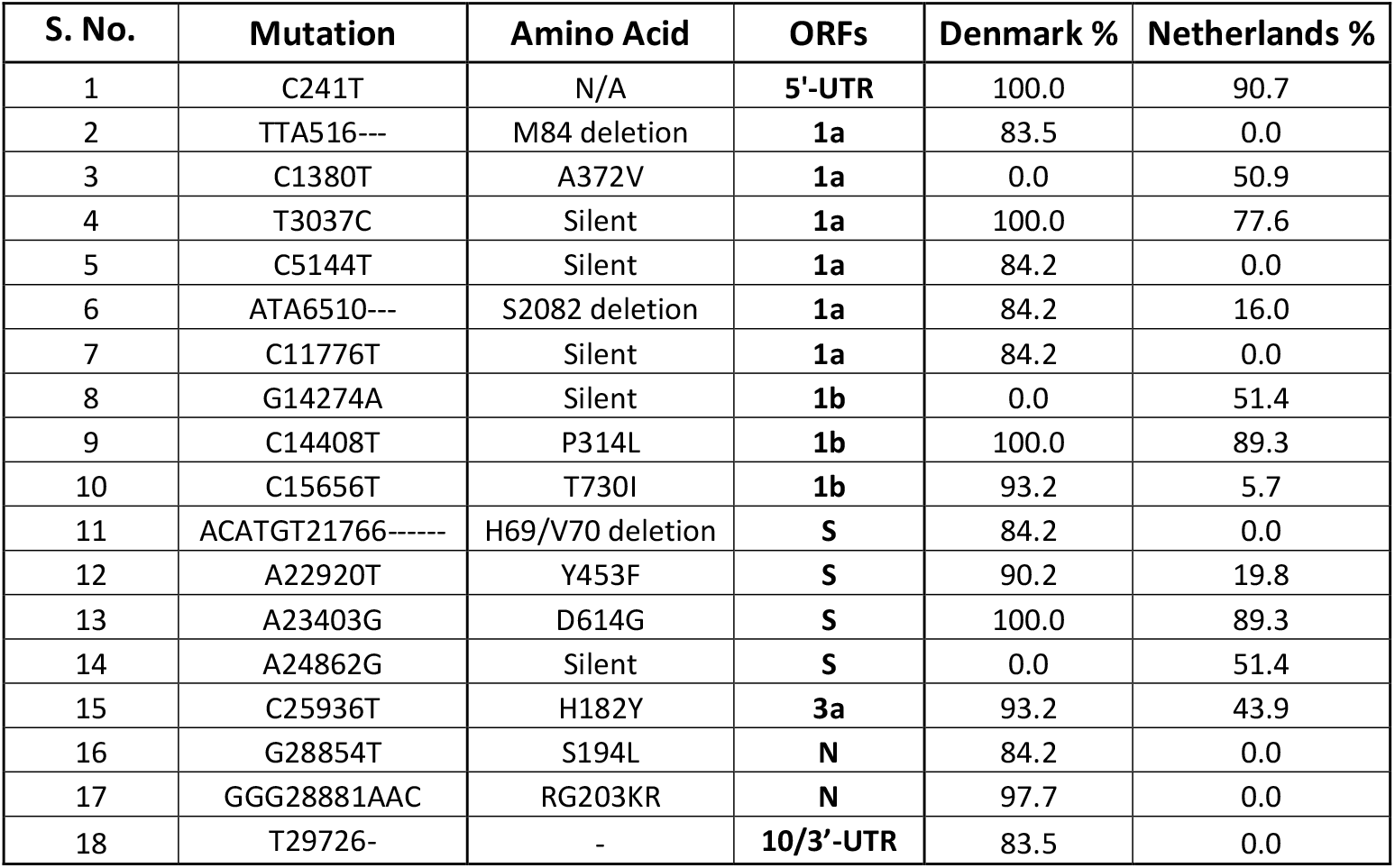
Mutations prevailed in mink derived SARS-CoV-2 genome.

### Mutations at Amino Acid Level

We translated all the 201 SARS-CoV-2 genomic mutations and found majority of the mutations (122) are non-synonymous mutations, which lead to amino acid changes while the ORF-M and 7a had more synonymous (4 out of 6 and 2 out of 3, respectively) than non-synonymous mutations, Figure 3, and Supplementary Table 1. The five mutations, which do not code for any proteins, laid between ORFs, viz. ATATTAGTTTTTCTGTTTGG26487 deletion (between ORF-3a and M), G27390A (between ORF-6 and 7a), CTTATT27697deletion (between ORF-7a and 7b), T29685C, and T29726 deletion (between ORF-10 and 3’-UTR), Supplementary Table 1.

**Figure 4:**
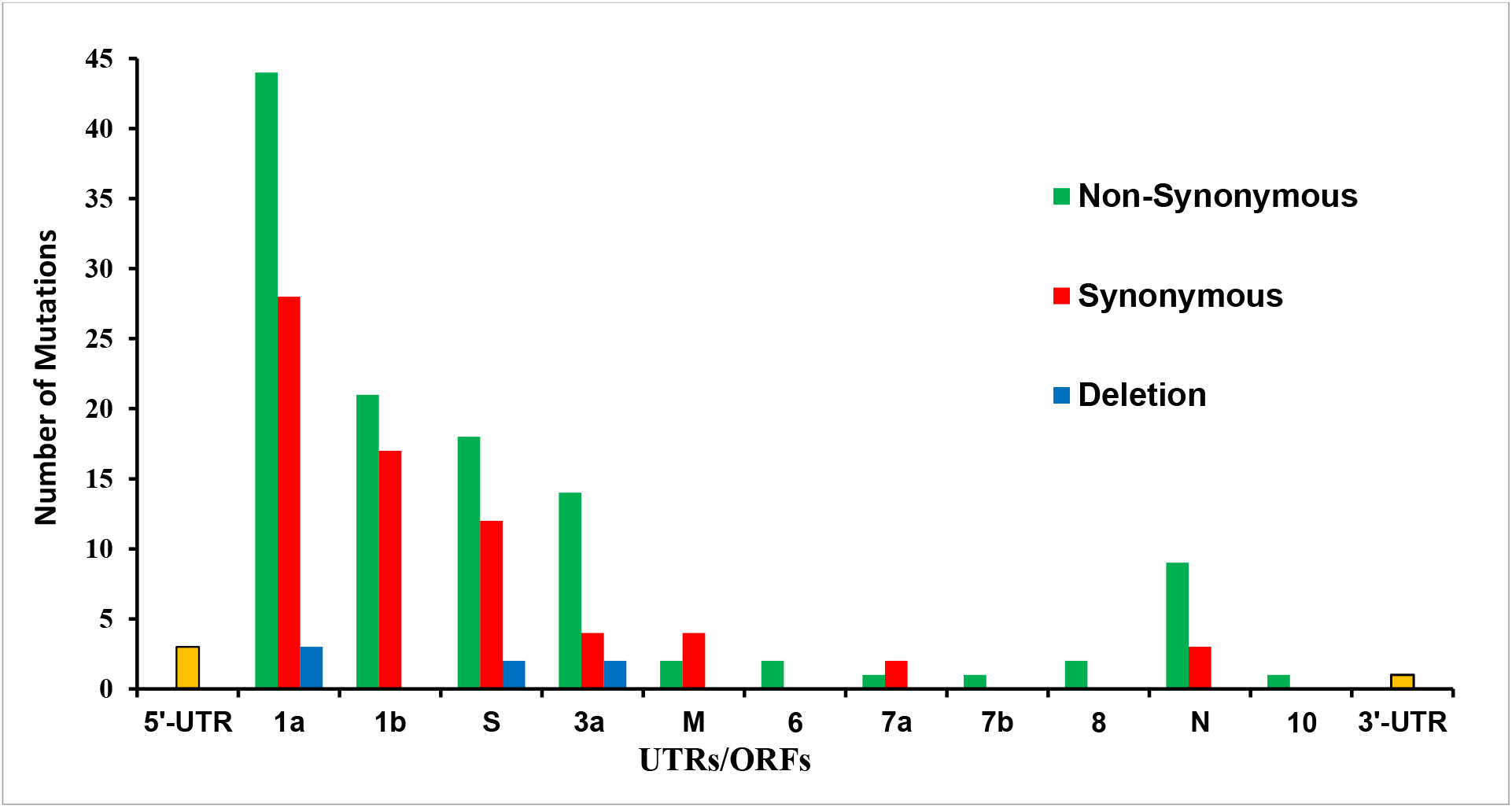
Mutations at amino acid level.

### The Frame-Shift and Non-Sense Mutations

During the mutational analysis, we found TTAATCCAGTA26159 deletion in 1.9% of Netherlands’ samples lead to frame-shift mutation, Supplementary Table 1. The mutation laid at position 256 of the ORF-3a, which is 276 amino acid long ORF, lead to change in amino acid sequences after the position 256. We looked the frame-shift mutation in the January 2021 set of data and we did not find any other sample with this mutation.

Another mutation G27948T in 2.3% of Denmark derived samples, which laid in the ORF-8 translated to stop codon after amino acid 18 in the ORF, which is 122 amino acid long. We also analysed non-sense mutation in the January 2021 set of data and we found 15 samples, all were derived from Denmark samples, had this mutation, Supplementary Table 2, which led the truncated ORF-8 expression.

## Discussions

The binding capability of the SARS-CoV-2 S protein to the ACE-2 receptor of a host’s cells makes the cells susceptible to the viral entry and that determines the hosts’ range and tissue tropism of the SARS-CoV-2. Table 1 reveals the broad mammalian hosts’ range of the SARS-CoV-2. However, for productive viral replication, mere viral entry is not enough; compatibilities of host’s cellular machinery are also essential, and that could be a reason for a handful of reports of the SARS-CoV-2 infection to domesticated animals, although it is obvious that many instances of exposure of the virus to the domesticated animals would have happened. There could be host factors that restrict the virus to replicate and transmit in those mammals. These plausible factors need to be explored for their therapeutic applications in human counterparts.

The first transmission of the virus to humans in the Wuhan seafood market was said to be from bat/pangolin (25), but we have only a few SARS-CoV-2 genomes derived from these suspected animals as shown in Table 1. Rather, the 755 genomic sequences derived from the mink (*Neovison vison*) and the majority of minks found infected in many farms in the Netherlands and Denmark (7–10), indicate the minks are highly susceptible to the viral infection. The phylogenetic tree shown in Figure 2, describes the similarities in the genomic sequences of the virus; the human-derived viral sequence is more similar with minks derived viral sequences than bat or pangolin derived viral sequences. The close resemblance of human and mink derived SARS-CoV-2 make the virus transmission to human and mink back and forth (8), which can make minks a potential SARS-CoV-2 reservoir.

This study finds 201 mutations from the analysis of 347 samples (214 from the Netherlands and 133 from Denmark) available in GISAID. The mutation patterns in these genomic sequences were very different (Supplementary Table 1). Out of the 201 mutations found, 113 were from Netherlands derived sample while 101were from Denmark derived samples. Only 13 mutations were found common in Netherlands’ and Denmark’s derived samples (Supplementary Table 1). From these 13 common mutations, 8 mutations (C241T, T3037C, ATA6510 deletion, C14408T, C15656T, A22920T, A23403G, and C25936T) prevailed in either Netherland or Denmark derived samples (Table 2 and Supplementary Table 1). Out of these 8, 4 mutations (C241T, T3037C, C14408T, and A23403G) prevailed in both Netherland and Denmark derived samples. Interestingly, these 4 mutations were also reported to prevail in human-derived SARS-CoV-2 (20). Of the 201 mutations we found, the majority of them (122) were non-synonymous mutations, studies on human-derived SARS-CoV-2 also revealed similar results (14–16, 18), suggesting SARS-CoV-2 proteins are capable of tolerating amino acid sequences change, which has the potential to question the success of available vaccines and upcoming drugs.

We have found TTAATCCAGTA26159 deletion in 4 samples derived from the Netherlands, which leads to frame-shift mutation after amino acid 255 in the ORF-3a, which is 276 amino acid long protein, Supplementary Table 1 and Supplementary Table 2. We analyzed the January 2021 set of data, but we haven’t found the deletion mutation in any sample except those four samples mentioned in Supplementary Table 2.

Another mutation G27948T in ORF-8, which generates a stop codon after 18 amino acids, found in two samples derived from Denmark. We analyzed the January 2021 set of data for the mutation and found 15 samples, all from Denmark, were had the non-sense mutation, Supplementary Table 1 and Supplementary Table 3. A similar mutation also reported from the Khalid et al study, where 17 amino acids long peptide were reported instead of 18 amino acids, which shows that the SARS-CoV-2 can replicate without the full-length ORF-8 protein.

In this study, all mutation analysis of the SARS-CoV-2 genomes, we performed in reference to Wuhan derived sequence (NC_22452.2), which establishes close resemblance in the gnome sequences derived from human and mink hosts. Although the mutation patterns are different in mink-derived samples from the Netherland and Denmark, different patterns have also been found in human-derived SARS-CoV-2 genome sequences (20). On the other hand, some of the common mutations are prevailing in the SARS-CoV-2 genomes, derived from human and mink hosts. These findings further validate the similarities of the SARS-CoV-2 genomes derived from human and mink hosts.

## Conclusions

The SARS-CoV-2 infection to humans is a zoonotic spillover. The spillover facilitated by interactions of viral proteins and the host’s cellular machinery. The mutation capability not only helps the virus to escape from vaccines and antiviral drugs but also provides unlimited possibilities to cross the species barriers. The susceptibility of minks to SARS-CoV-2 infection is another example of zoonotic spillover. We do not have reasons to believe that the zoonotic spillover would not happen again. It is just a matter of time when mutant variants of the virus will find compatible hosts. As reports suggest (8), the virus transmitted human and mink back and forth, and that would also be possible to any new host, the virus may find.

## Acknowledgment

We would like to extend our thanks to the people at NCBI database and GISAID for making the SARS-CoV-2 genome sequences available.

## Author Contributions

Conception/design: MK

Data collection: MK

Data analysis/interpretation: MK, YA

Drafting article: MK, YA

Critical revision of the article: MK, YA

Verifying underlying data: MK, YA

## Abbreviations

GISAID: Global initiative on sharing all influenza data
NCBI: National Center for Biotechnology Information
NSP6: Non-structural protein 6
ORF-8: Open Reading Frame-8
UTRs: Untranslated regions

## References

1. Mattia Mori, Clemente Capasso, Fabrizio Carta, William a Donald & Claudiu T Supuran (2020) A deadly spillover: SARS-CoV-2 outbreak, Expert Opinion on Therapeutic Patents, 30:7, 481–485, DOI: 10.1080/13543776.2020.1760838

2. SA Kemp, RP Datir, DA Collier, IATM Ferreira, A Carabelli, W Harvey, DL Robertson, and RK Gupta. Recurrent emergence and transmission of a SARS-CoV-2 Spike deletion ΔH69/V70. bioRxiv. DOI: https://doi.org/10.1101/2020.12.14.422555

3. https://www.bbc.com/news/health-55388846

4. Wang Q, Zhang Y, Wu L, Niu S, Song C, Zhang Z, Lu G, Qiao C, Hu Y, Yuen KY, Wang Q, Zhou H, Yan J, Qi J. Structural and functional basis of SARS-CoV-2 entry by using human ACE2. Cell 181, 1–11 (2020). DOI: 10.1016/j.cell.2020.03.045

5. Lan J, Ge J, Yu J, Shan S, Zhou H, Fan S, Zhang Q, Shi X, Wang Q, Zhang L, Wang Z. Structure of the SARS-CoV-2 spike receptor-binding domain bound to the ACE2 receptor. Nature 581, 215–220 (2020). https://doi.org/10.1038/s41586-020-2180-5

6. Shang J, Ye G, Shi K, Wan Y, Luo C, Aihara H, Geng Q, Auerbach A, Li F. Structural basis of receptor recognition by SARS-CoV-2. Nature 581, 221–224 (2020). DOI: https://doi.org/10.1038/s41586-020-2179-y

7. Oreshkova N, Molenaar RJ, Vreman S, Harders F, Oude Munnink BB, Hakze-van der Honing RW, Gerhards N, Tolsma P, Bouwstra R, Sikkema RS, Tacken MG, de Rooij MM, Weesendorp E, Engelsma MY, Bruschke CJ, Smit LA, Koopmans M, van der Poel WH, Stegeman A. SARS-CoV-2 infection in farmed minks, the Netherlands, April and May 2020. Euro Surveill. 2020 Jun;25(23):2001005. DOI: 10.2807/1560-7917.

8. Oude Munnink BB, Sikkema RS, Nieuwenhuijse DF, Molenaar RJ, Munger E, Molenkamp R, van der Spek A, Tolsma P, Rietveld A, Brouwer M, Bouwmeester-Vincken N, Harders F, Hakze-van der Honing R, Wegdam-Blans MCA, Bouwstra RJ, GeurtsvanKessel C, van der Eijk AA, Velkers FC, Smit LAM, Stegeman A, van der Poel WHM, Koopmans MPG. Transmission of SARS-CoV-2 on mink farms between humans and mink and back to humans. Science. 2021 Jan 8;371(6525):172–177. DOI: 10.1126/science.abe5901

9. Enserink M. Coronavirus rips through Dutch mink farms, triggering culls. Science. 2020 Jun 12;368(6496):1169. DOI: 10.1126/science.368.6496.1169

10. Hobbs EC, Reid TJ. Animals and SARS-CoV-2: Species susceptibility and viral transmission in experimental and natural conditions, and the potential implications for community transmission. Transbound Emerg Dis. 2020 Oct 22. DOI: 10.1111/tbed.13885

11. Toby S. Mink Infected Two Humans with Coronavirus: Dutch Government. Reuters. (2020) Available online at: https://www.reuters.com/article/us-h (accessed October 19, 2020)

12. http://animalguardians.us/issues/fur-industry

13. Chan JF, Kok KH, Zhu Z, Chu H, To KK, Yuan S, Yuen KY. Genomic characterization of the 2019 novel human-pathogenic coronavirus isolated from a patient with atypical pneumonia after visiting Wuhan. Emerg Microbes Infect. 2020 Jan 28;9(1):221–236. doi: 10.1080/22221751.2020.1719902.

14. Peck KM, Lauring AS. 2018. Complexities of viral mutation rates. J Virol 92:e01031–17. DOI: https://doi.org/10.1128/JVI.01031-17

15. Pachetti M, Marini B, Benedetti F, Giudici F, Mauro E, Storici P, Masciovecchio C, Angeletti S, Ciccozzi M, Gallo R, Zella D, Ippodrino R. Emerging SARS-CoV-2 mutation hot spots include a novel RNA-dependent-RNA polymerase variant. J Transl Med 18, 179 (2020). DOI: https://doi.org/10.1186/s12967-020-02344-6

16. Phan T. Genetic diversity and evolution of SARS-CoV-2. Infection, Genetics and Evolution 81 (2020) 104260. DOI: https://doi.org/10.1016/j.meegid.2020.104260

17. Tang X, Wu C, Li X, Song Y, Yao X, Wu X, Duan Y, Zhang H, Wang Y, Qian Z, Cui J, Lu J. On the origin and continuing evolution of SARS-CoV-2. National Sci Rev. 2020. DOI: https://doi.org/10.1093/nsr/nwaa036

18. Wang C, Liu Z, Chen Z, Huang X, Xu M, He T, Zhang Z. The establishment of reference sequence for SARS-CoV-2 and variation analysis. J Med Virol. 2020 Jun;92(6):667–674. DOI: 10.1002/jmv.25762

19. https://www.gisaid.org

20. Khalid M, Murphy D, Shoai M, Al-ebini Y. Host’s Specific SARS-CoV-2 Mutations: Insertion of the Phenylalanine in the NSP6 Linked to the United Kingdom and Premature Termination of the ORF-8 Associated with the European and the United States of America Derived Samples. biorxiv, https://www.biorxiv.org/content/10.1101/2020.12.29.424530v2

21. http://multalin.toulouse.inra.fr/multalin/

22. Wu F, Zhao S, Yu B, Chen YM, Wang W, Song ZG, Hu Y, Tao ZW, Tian JH, Pei YY, Yuan ML, Zhang YL, Dai FH, Liu Y, Wang QM, Zheng JJ, Xu L, Holmes EC, Zhang YZ. Author Correction: A new coronavirus associated with human respiratory disease in China. Nature. 2020 Apr;580(7803):E7. DOI: 10.1038/s41586-020-2202-3

23. Kumar S, Stecher G, Li M, Knyaz C, Tamura K. MEGA X: Molecular Evolutionary Genetics Analysis across Computing Platforms. Mol Biol Evol. 2018 Jun 1;35(6):1547–1549. doi: 10.1093/molbev/msy096.

24. Korber B, Fischer WM, Gnanakaran S, Yoon H, Theiler J, Abfalterer W, Hengartner N, Giorgi EE, Bhattacharya T, Foley B, Hastie KM, Parker MD, Partridge DG, Evans CM, Freeman TM, de Silva TI; Sheffield COVID-19 Genomics Group, McDanal C, Perez LG, Tang H, Moon-Walker A, Whelan SP, LaBranche CC, Saphire EO, Montefiori DC. Tracking Changes in SARS-CoV-2 Spike: Evidence that D614G Increases Infectivity of the COVID-19 Virus. Cell. 2020 Aug 20;182(4):812–827.e19. DOI: 10.1016/j.cell.2020.06.043

25. Jo WK, de Oliveira-Filho EF, Rasche A, Greenwood AD, Osterrieder K, Drexler JF. Potential zoonotic sources of SARS-CoV-2 infections. Transbound Emerg Dis. 2021 Jul;68(4):1824–1834. doi: 10.1111/tbed.13872

